# A High-Resolution Genetic Map for the Laboratory Rat

**DOI:** 10.1101/268227

**Authors:** John Littrell, Shirng-Wern Tsaih, Amelie Baud, Pasi Rastas, Leah Solberg-Woods, Michael J Flister

## Abstract

An accurate and high-resolution genetic map is critical for mapping complex traits, yet the resolution of the current rat genetic map is far lower than human and mouse, and has not been updated since the original Jensen-Seaman map in 2004. For the first time, we have refined the rat genetic map to sub-centimorgan (cM) resolution (<0.02 cM) by using 95,769 genetic markers and 870 informative meioses from a cohort of 528 heterogeneous stock (HS) rats. Global recombination rates in the revised sex-averaged map (0.66 cM/Mb) did not difeer compared to the historical map (0.65 cM/Mb); however, substantial refinement was made to the localization of highly recombinant regions within the revised map. Also for the first time, sex-specific rat genetic maps were generated, which revealed both genomewide and fine-scale variation in recombination rates between male and female rats. Reanalysis of multiple quantitative trait loci (QTL) using the historical and refined rat genetic maps demonstrated marked changes to QTL localization, shape, and effect size. As a resource to the rat research community, we have provided revised centimorgan positions for all physical positions within the rat genome and commonly used genetic markers for trait mapping, including 44,828 SSLP markers and the RATDIV genotyping array. Collectively, this study provides a substantial improvement to the rat genetic map and an unprecedented resource for analysis of complex traits and recombination in the rat.

## INTRODUCTION

Genetic maps are constructed by calculating the frequency of meiotic recombination between genomic markers to define the linear order and relative distance between loci. In addition to refining the physical map of a species (i.e., the genomic sequence), a genetic map is fundamentally important for the analysis of quantitative trait loci (QTL) that link disease phenotypes to genomic intervals containing disease modifiers. The accuracy of a genetic map influences the correct calculation of a QTL intervals, because the linkage of a trait to a genomic interval is a function of the genetic distance (i.e., the expected recombination frequency) between the two flanking markers (LANDER and BOTSTEIN 1986). Thus, miscalculating the genetic distance between two flanking markers is expected to alter the QTL interval, which ultimately impacts the list of candidate disease modifier(s) within the QTL. Indeed, a recent refinement to the mouse genetic map revealed dramatic changes to previously characterized QTL (COX et al. 2009).

One way to improve the resolution of a genetic map is to increase the number of meiotic events and genetic markers that are assessed. This approach has been adopted for constructing high resolution genetic maps of the human (104,246 meioses; 833,754 markers) (BHERER et al. 2017) and mouse (15,832 meioses; 120,789 markers) (MORGAN et al. 2017). However, in stark contrast, the genetic map for the rat has never been updated and is currently based on a considerably smaller dataset (90 meioses; 2,305 markers) (JENSEN-SEAMAN et al. 2004; STEEN et al. 1999), with a map resolution of only 1.1 cM, as compared with resolutions of <0.004 cM (BHERER et al. 2017) and <0.01 cM(MORGAN et al. 2017) for the human and mouse genetic maps, respectively. As such, the relatively low resolution of the current rat genetic map is a potential limitation for accurately mapping QTL in the rat.

Here, we generated the first high resolution sex-averaged genetic map of the rat using 95,769 informative markers and 528 NIH-Heterogeneous Stock (HS) rats (65 families; 870 meioses), with an average resolution of <0.02 cM. Additionally, we produced the first sex-specific genetic maps for the rat, which recapitulated the dimorphic recombination rates observed in human (BHERER et al. 2017) and mouse (COX et al. 2009). Finally, reanalysis of historical QTL using the refined rat genetic maps revealed changes to QTL localization, shape, and effect size.

## MATERIALS AND METHODS

### Rats

The HS rat colony was initiated by the National Institutes of Health in 1984 using eight inbred strains (ACI/N, BN/SsN, BUF/N, F344/N, M520/N, MR/N, WKY/N, and WN/N) (HANSEN and SPUHLER 1984).

### Genotyping and Data Cleaning

The collection of genotypes from 528 HS rats was described previously (BAUD et al. 2014; RAT GENOME et al. 2013). Briefly, genomic DNA were fragmented with NspI and StyI endonucleases and hybridized to the custom RATDIV array that assays 797,968 genotypes spread evenly across the rat genome, with a median inter-SNP distance of 3,470bp (RGSC6.0/rn6). High quality genotypes were called using the BRLMM-P component of the Apt-Probeset-Genotype program that is part of the Affymetrix Power Tools suite (apt-1.10.2), as described previously (RAT GENOME et al. 2013). The genotype data were further cleaned to remove monomorphic SNPs and genotypes with high Mendelian inheritance rate were identified by PLINK and removed (PURCELL et al. 2007). After data cleaning, a total of 412,430 high-quality polymorphic genotypes remained, which far exceeds the number of genotype markers required for accurate estimation of the rat genetic map. Thus, we performed an unbiased selection of one genotype (MAF > 0.05) per every 10Kb window across the genome, which yielded a final list of 132,521 informative genotypes that were uniformly distributed across the genome.

### Construction of the High-Resolution Rat Genetic Map

The rat genetic map was constructed using Lep-MAP3 (LM3) (RASTAS 2017), which is a linkage mapping program designed for large genotyping datasets, such as the high-density genotyping data used in our study. The following LM3 functions were used to construct the rat genetic map for each of the 20 autosomes and for the X chromosome: 1) the ParentCall2 function was used to call parental genotypes by taking into account genotype information of grandparents, parents, and offspring; 2) the Filtering2 function was used to remove those markers with segregation distortion or those with large amounts of missing data; and 3) the OrderMarkers2 function was used to compute cM distances (i.e. recombination rates) between all adjacent markers per chromosome. Although LM3 was previously shown to be highly accurate in ordering markers (RASTAS 2017), we detected multiple markers with cM positions in obvious disagreement with their reported physical position in the RGSC 6.0/rn6 genome assembly. As this could be due to multiple factors and would potentially distort the genetic map, we sought to systematically remove errant markers with high disagreement between genetic and physical positions. A two-stage process was used for map cleaning a given chromosome for each of the sex-specific genetic distances produced by LM3 to build sex-specific and sex-averaged genetic maps. First, a locally weighted regression model (LOESS) (CLEVELAND and DEVLIN 1988) was used to assess the relationship between physical (bp) and genetic (cM) positions. Markers with absolute residuals of >1 were identified and removed since they disrupt the expected monotonically increasing behavior between the physical and genetic maps. Second, a cubic splines model was fitted to the plot of genetic versus physical distances of the remaining markers to estimate local recombination rates and to interpolate the genetic positions of markers on the original SNP array for each gender, based on their positions in RGSC 6.0/rn6. A final set of 95,769 informative markers were analyzed across ten independent replicate LM3 analyses and were used to calculate the sex-averaged and sex-specific rat genetic maps.

### Assigning Recombination Rates to Physical Positions and Genotyping Markers

Sex-averaged and sex-specific centimorgan distances were assigned to every kilobase of the physical map (RGSC 6.0/rn6) by interpolation with the revised rat genetic maps.Likewise, sex-averaged and sex-specific centimorgan distances were assigned to each genotype position within the RATDIV array (RAT GENOME et al. 2013). A total of 44,828 SSLP markers with physical positions were also given sex-specific and sex-averaged centimorgan distances by interpolation with the revised rat genetic maps.

### QTL Reanalysis

To assess the impact of the revised rat genetic map on QTL mapping, we re-examined the data from a previously published F2 reciprocal cross using WKY and F344 rats (SOLBERG et al. 2004). The Animal Care and Use Committee approval of the study and a detailed description of the methods can be found in the original article (SOLBERG et al. 2004). Data were downloaded directly from the Mouse Phenome Database (MPD; https://phenome.jax.org/centers/QTLA) and QTL were reanalyzed using the R/qtl software package (BROMAN et al. 2003) (version 1.41-6; http://www.rqtl.org/). The primary phenotypes-of-interest were related to depression-like behavior in rodents (e.g., immobility and climbing). Four F2 rats with missing data on both immobility and climbing (3 male and 1 female) and four SSLP markers with a high rate of missing data (D6Rat24, D6Rat128, D6Rat88, D6Mgh4) were excluded. A total of 482 F2 rats (222 females and 260 males) and 101 autosomal SSLP markers were used in the QTL re-analysis. No other phenotypic outliers or problematic markers were detected, as would be expected from using pre-cleaned data from the MPD. A single-locus genomewide QTL scan was performed and LOD scores were calculated at 2 cM interval across the genome, using the imputation method implemented in R/qtl (SEN and CHURCHILL 2001) and significance was determined on the basis of 1000 permutations of the data (CHURCHILL and DOERGE 1994). A LOD score exceeding the 0.05 genomewide adjusted threshold was considered significant, whereas a LOD score exceeding the 0.63 genomewide adjusted threshold was considered to be suggestive (LANDER and KRUGLYAK 1995). The Bayes credible interval function in R/qtl (bayesint) was used to approximate the 95% confidence intervals for the QTL peak location for both the additive and the interactive models, as described in (SOLBERG et al. 2004). QTL that were detected using the original or repositioned markers were compared for correspondence in peak positioning, shape, and significance level, as described previously (COX et al. 2009).

### Data Availability

File S1 contains detailed descriptions of all supplemental files. File S2 contains sex-averaged and sex-specific centimorgan distances that were assigned to every kilobase of the physical map (RGSC 6.0/rn6) by interpolation with the revised rat genetic maps. File S3 contains sex-averaged and sex-specific centimorgan distances for each genotype position within the RATDIV array. File S4 contains sex-averaged and sex-specific centimorgan distances for 44,828 SSLP markers with assigned physical positions from the Rat Genome Database (http://rgd.mcw.edu/)

**Figure 1.**
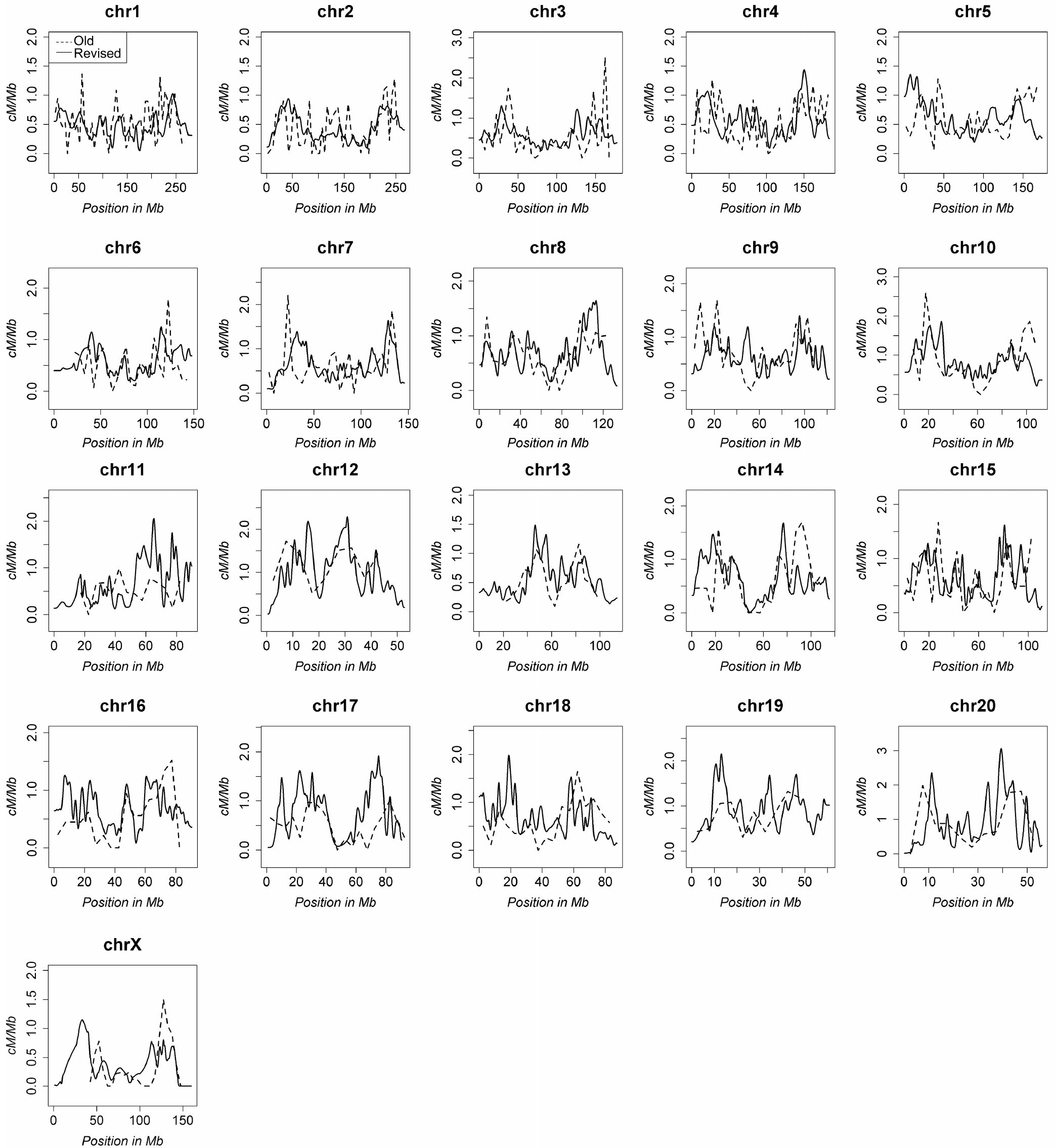
Comparison of the original and revised sex-averaged rat genetic maps. The recombination rates for all chromosomes are compared between the original Jensen-Seaman map (hashed line) and the revised rat genetic map (solid line).

## RESULTS

### Construction of the high-resolution rat genetic map

To generate the first high-density genetic map of the laboratory rat, genomewide genotyping data were collected from 528 HS rats using the RATDIV high-density SNP array. Following data cleaning (see methods section), a final set of 95,769 informative markers were used to calculate the sex-specific centimorgan distances using the LM3 program(RASTAS 2017). The average resolution of the revised rat genetic map was <0.02 cM, which is considerably higher than the 1.1 cM resolution of the previous map (JENSEN-SEAMAN et al. 2004). As shown in Table 1, the sizes of the most recent physical map (RGSC 6.0/rn6) and the revised genetic map were both increased by roughly 10% compared with the circa 2004 assemblies of the rat physical map (Baylor 3.4/rn4) and rat genetic map (JENSEN-SEAMAN et al. 2004), yielding comparable average recombination rates of 0.66 cM/Mb and 0.65 cM/Mb in the Jensen-Seaman map and the revised rat genetic map, respectively. Compared with males, the female-specific rat genetic map was approximately 10% larger (1,826 cM vs. 1,589 cM), which is similar to the dimorphic recombination reported in mouse (COX et al. 2009) and human (BHERER et al. 2017). To our knowledge, a sex-specific genetic map for the rat has not previously been reported, precluding the comparison of sex-specific recombination rates with historical data. Nonetheless, these data suggest that the revised genetic map is of comparable size to the Jensen-Seaman map, when accounting for the larger physical assembly of RGSC 6.0/rn6 compared with the Baylor 3.4/rn4 genome assembly. Moreover, the revised rat genetic map provides novel insight to the disparate recombination rates between males and females.

### Updated distribution of recombination rates in the rat genome

Although the average recombination rates of the rat genome did not differ greatly from the Jensen-Seaman map, we observed marked shifts in the distribution and amplitudeof recombination frequencies in the revised genetic map (Figure 1). We attribute the altered distribution of recombination rates within the revised genetic map to more accurate marker placement, which is likely due to the greater number of informative haplotypes within the HS population, as well as a 10-fold increase in informative meioses and >20-fold higher map resolution compared with the historical Jensen-Seaman map. Another noted advantage of the revised rat genetic map is the characterization of dimorphic recombination rates, which to our knowledge has not previously been reported for the rat. As found in other mammalian genomes(BHERER et al. 2017; COX et al. 2009), a substantial portion of the rat genome showed dimorphic recombination rates and higher recombination rates were typically observed in females (Figure 2). However, despite the genomewide trend of increased recombination in females compared with males, fine-scale variation in dimorphic recombination rates was pervasive and highly recombinant regions in both sexes were detected. Collectively, these data reveal for the first time the dimorphic recombination rates in the rat.

### Impact of the revised rat genetic map on QTL identificatio

Accurate marker placement is crucial for correctly localizing QTL. To assess whether the revised rat genetic map affects QTL localization, we repeated the linkage analysis from a previously reported reciprocal F2 cross of WKY and Fisher 344 rats (SOLBERG et al. 2004). Identical methodology was used to calculate QTL, except for altering the marker positions based on the Jensen-Seaman map or the revised rat genetic map. Phenotypic data (e.g., immobility and climbing scores) from the F2 generation male and female rats (total n=486) were retrieved from the MPD and QTL were calculated using historical and revised marker positions. Reanalysis of the QTL using the original sex-averaged marker positions from the Jensen-Seaman map recapitulated all 12 QTL that were previously reported for rat immobility and climbing phenotypes in the F2 generation. In comparison, analysis of the QTL using the revised genetic map revealed multiple QTL with altered shape, localization, and effect size. A comparison of the peak LOD scores and 95% confidence intervals (CI) of QTL calculated with the original or revised maps revealed 9 out of 32 QTL with shifts of greater than 10 cM (Table 2). Collectively, these data demonstrate that the accuracy of marker placement has a dramatic impact on QTL localization, which would subsequently impact identification of the underlying causative gene(s).

### Assignment of centimorgan distances to the physical map and all SSLP markers within the rat genome

As a resource to the scientific community, we have provided the sex-averaged and sex-specific centimorgan distances per every kilobase of the current physical map (RGSC 6.0/rn6) (File S2). We have also provided the revised centimorgan positions for every genotype position within the RATDIV genotyping array (RAT GENOME et al. 2013) (File S3) and for 44,828 rat SSLP markers with unique physical positions that are currently annotated in the Rat Genome Database (rgd.mcw.edu) (File S4). Finally, in the process of curating the known SSLP markers, we identified 8,457 markers in the RGD database that aligned to multiple positions or were assigned no position in the current physical genome build and we therefore do not recommend using these SSLP markers for future mapping studies.

## DISCUSSION

In contrast to frequent revisions of high-resolution human and mouse genetic maps (BHERER et al. 2017; COX et al. 2009), the rat genetic map has been restricted to a relatively low-resolution (1.1 cM) map for the past two decades (JENSEN-SEAMAN et al. 2004). To address this issue, we constructed a revised rat genetic map from a single large cohort of 528 HS rats, which is to our knowledge the most diverse genetic resource for map building in the rat (capturing haplotypes from 8 parental strains) and the highest-resolution map (<0.02 cM) of any rat genetic map to date. These data were used to assign highly accurate centimorgan positions, for the first time, to all physical positions within the rat genome (RGSC 6.0/rn6), to all genotypes captured by the RATDIV array, and to 44,828 SSLP markers by interpolation with a high-resolution, revised rat genetic map. To our knowledge, this is also the first rat genetic map to analyze both sex-averaged and sex-specific recombination rates, which were used to generate sex-specific rat genetic maps for the first time. Using these data, we demonstrate that QTL localization is affected by the accuracy of marker positions. Collectively, the current study provides a new consensus rat genetic map and a common framework for candidate gene mapping in the rat.

**Figure 2.**
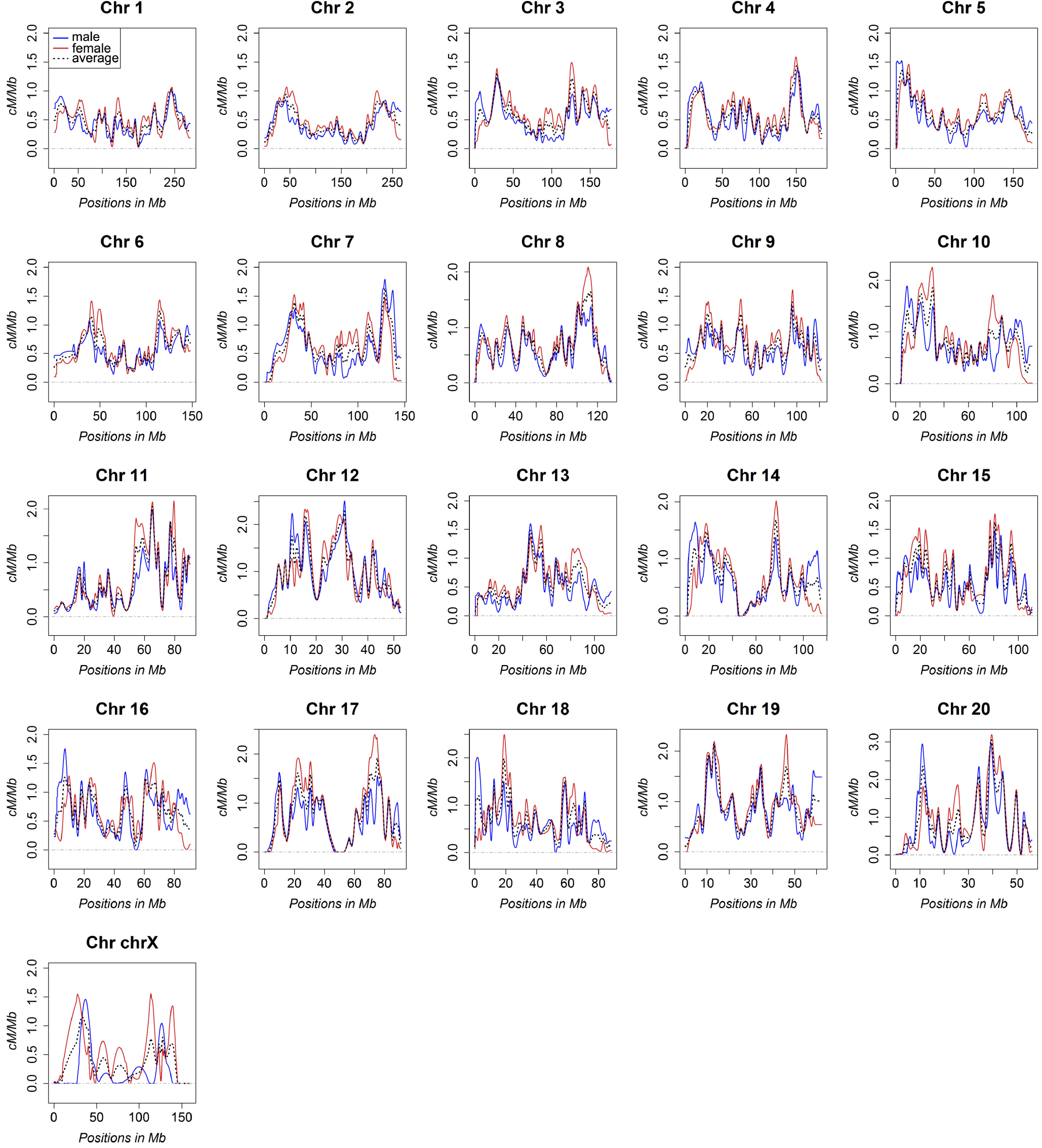
Comparison of the sex-specific rat genetic maps. The recombination rates for all chromosomes are compared between the male (blue) and female (red) rat genetic maps. For reference, the revised sex-averaged map is also plotted (black hashed line)

**Table 1.** Genetic and physical map of rat chromosomes and chromosome wide recombination rate

|  | Number<br>of Markers | Male<br>(cM) | Female<br>(cM) | Average<br>(cM) | Physical<br>Size (Mb) | Recombination<br>Rate (cM/Mb) | Average<br>(cM) | Physical<br>Size (Mb) | Recombination<br>Rate (cM/Mb) |
| --- | --- | --- | --- | --- | --- | --- | --- | --- | --- |
| chr1 | 9,098 | 134.4 | 146.0 | 140.2 | 282.8 | 0.50 | 149.3 | 267.4 | 0.56 |
| chr2 | 10,440 | 122.0 | 135.5 | 128.8 | 266.4 | 0.48 | 112.4 | 255.3 | 0.44 |
| chr3 | 6,142 | 95.4 | 113.1 | 104.2 | 177.7 | 0.59 | 91.5 | 166.0 | 0.55 |
| chr4 | 6,930 | 100.5 | 110.7 | 105.6 | 184.2 | 0.57 | 102.2 | 186.1 | 0.55 |
| chr5 | 6,859 | 99.3 | 111.8 | 105.6 | 173.7 | 0.61 | 105.7 | 171.6 | 0.62 |
| chr6 | 5,747 | 81.4 | 94.9 | 88.1 | 148.0 | 0.60 | 85.2 | 134.2 | 0.63 |
| chr7 | 5,157 | 91.6 | 99.4 | 95.5 | 145.7 | 0.66 | 88.5 | 141.6 | 0.62 |
| chr8 | 4,759 | 83.0 | 99.6 | 91.3 | 133.3 | 0.68 | 84.1 | 126.8 | 0.66 |
| chr9 | 3,761 | 72.8 | 83.9 | 78.3 | 122.1 | 0.64 | 78.3 | 109.5 | 0.71 |
| chr10 | 4,479 | 88.0 | 94.3 | 91.1 | 112.6 | 0.81 | 92.8 | 101.2 | 0.92 |
| chr11 | 2,788 | 56.0 | 66.8 | 61.4 | 90.5 | 0.68 | 39.0 | 73.6 | 0.53 |
| chr12 | 2,108 | 53.9 | 52.4 | 53.2 | 52.7 | 1.01 | 51.9 | 43.6 | 1.19 |
| chr13 | 4,869 | 59.6 | 70.3 | 64.9 | 114.0 | 0.57 | 43.9 | 75.1 | 0.58 |
| chr14 | 4,519 | 72.3 | 74.1 | 73.2 | 115.5 | 0.63 | 70.9 | 105.7 | 0.67 |
| chr15 | 2,991 | 64.4 | 79.2 | 71.8 | 111.2 | 0.65 | 63.8 | 106.1 | 0.60 |
| chr16 | 3,806 | 62.5 | 61.0 | 61.8 | 90.7 | 0.68 | 45.5 | 76.6 | 0.59 |
| chr17 | 2,679 | 61.2 | 74.4 | 67.8 | 90.8 | 0.75 | 43.9 | 91.0 | 0.48 |
| chr18 | 3,223 | 55.3 | 63.1 | 59.2 | 88.2 | 0.67 | 52.3 | 84.7 | 0.62 |
| chr19 | 2,360 | 51.2 | 58.5 | 54.8 | 62.3 | 0.88 | 48.1 | 56.4 | 0.85 |
| chr20 | 1,890 | 47.2 | 54.7 | 51.0 | 56.2 | 0.91 | 48.2 | 50.6 | 0.95 |
| chrX | 1,165 | 37.1 | 82.8 | 59.9 | 160.0 | 0.37 | 44.6 | 130.8 | 0.34 |
| Total | 95,769 | 1,589 | 1,826 | 1,708 | 2,619 | 0.66 | 1,542 | 2,554 | 0.65 |

### Changes to the rat genetic map and recombination rates

After accounting for the larger physical size of the RGSC 6.0/rn6 rat genome build (2,619 Mb) compared with the original Baylor 3.4/rn4 rate genome build (2,554 Mb), the increased size of the rat genetic map (1,708 cM) is proportional to the original Jensen-Seaman map (1,542 cM). Thus, although the coordinates of highly recombinant regions in the rat genome were refined in the revised rat genetic map, the sex-averaged genomewide recombination rates did not change (0.66 cM/Mb vs. 0.65 cM/Mb). Although the genomewide recombination rates did not change, fine-scale localization of highly recombinant regions differed between the Jensen-Seaman map and the revised rat genetic map. One potential reason for the refined localization of highly recombinant regions in the revised rat genetic map is the greater potential of genetic variation due to the possibility of eight informative HS founder haplotypes per genomic position, whereas prior rat genetic maps relied on crosses between two parental strains with less genetic variation (BIHOREAU et al. 2001; JENSEN-SEAMAN et al. 2004). In some instances within the Jensen-Seaman map, recombinations likely occurred, but were poorly localized due to sparse marker density and the limited number of informative meioses.

Thus, it stands to reason that the increased genetic diversity of our cohort of 528 HS rats, combined with the greater number of informative meioses (870 vs 90), and the higher density of genetic markers (95,769 vs 2,305), would enable more precise localization of crossover events in the revised map compared with the Jensen-Seaman map that was based on a much smaller cross (45 rats) of only two strains (SHRSP x BN).

In addition to improving map resolution, we have analyzed the first sex-specific genetic maps for the rat. Similar to human (BHERER et al. 2017) and mouse (COX et al. 2009), the female-specific rat genetic map was larger than the male-specific map, indicating that the genomewide average of recombination events are more frequent in female rats compared with males. Nonetheless, fine-scale variability in the ratio of male-to-female recombination is pervasive across the genome and multiple highly recombinant regions that are unique to either gender were detected. Notably, the underlying mechanisms of dimorphic recombination have yet to be examined in the rat, as sex-specific genetic maps were not previously available. Several mechanisms of dimorphic recombination have been proposed in other species though, including the sex-specific crossover interference (PETKOV et al. 2007), sex-specific molecular mediators of recombination (BHERER et al. 2017), and sex-specific haploid selection (LENORMAND and DUTHEIL 2005). The current findings suggest that similar sex-specific mechanisms potentially underlie the dimorphic recombination rates in the rat and could be explored using the HS model.

**Table 2.**
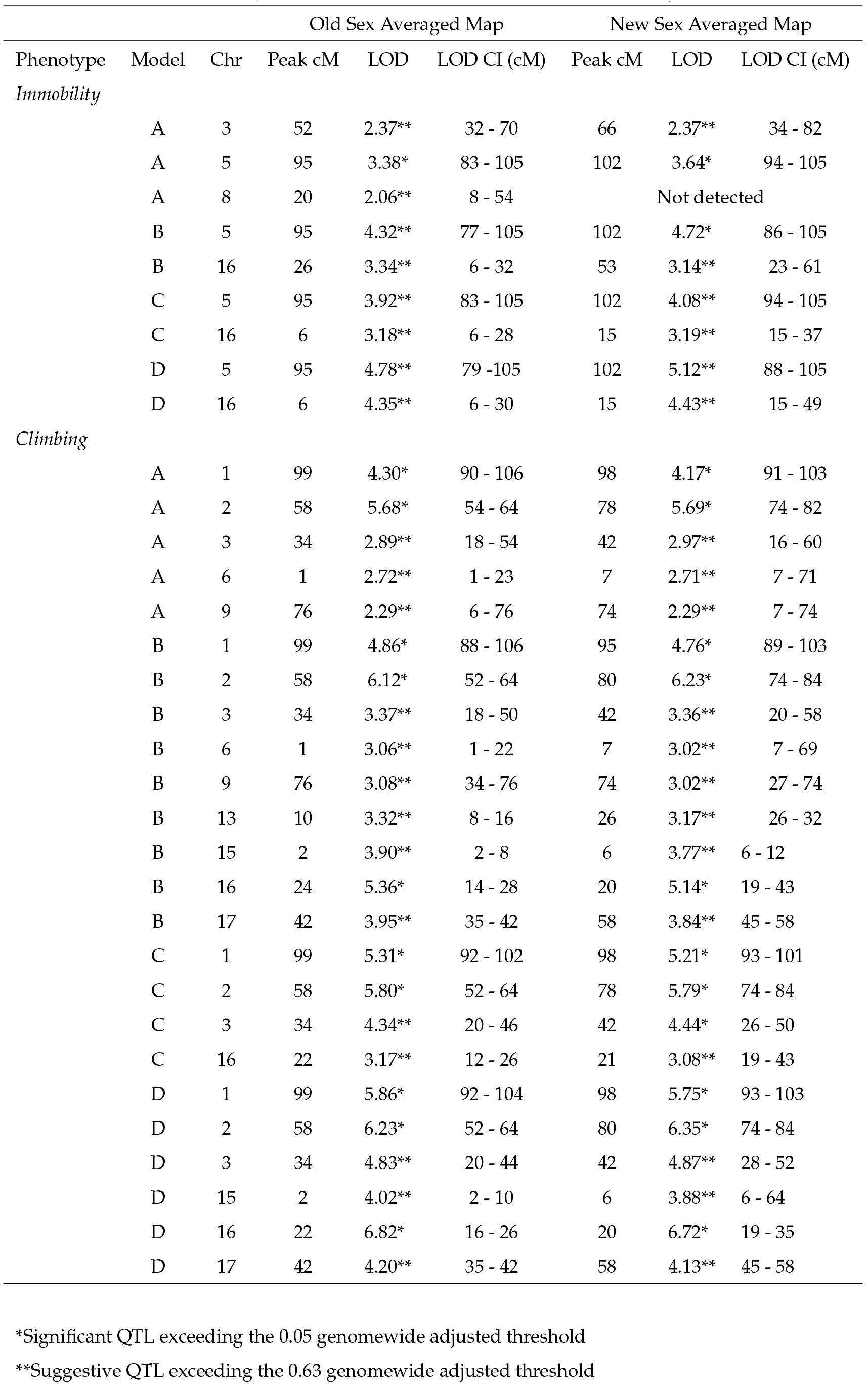
Comparison of QTL calculated using the old and revised sex-averaged rat genetic maps

| Phenotype | Model | Chr | Old Sex Averaged Map |  |  | New Sex Averaged Map |  |  |
| --- | --- | --- | --- | --- | --- | --- | --- | --- |
|  |  |  | Peak cM | LOD | LOD CI (cM) | Peak cM | LOD | LOD CI (cM) |
| *Immobility* |  |  |  |  |  |  |  |  |
|  | A | 3 | 52 | 2.37** | 32 - 70 | 66 | 2.37** | 34 - 82 |
|  | A | 5 | 95 | 3.38* | 83 - 105 | 102 | 3.64* | 94 - 105 |
|  | A | 8 | 20 | 2.06** | 8 - 54 | Not detected |  |  |
|  | B | 5 | 95 | 4.32** | 77 - 105 | 102 | 4.72* | 86 - 105 |
|  | B | 16 | 26 | 3.34** | 6 - 32 | 53 | 3.14** | 23 - 61 |
|  | C | 5 | 95 | 3.92** | 83 - 105 | 102 | 4.08** | 94 - 105 |
|  | C | 16 | 6 | 3.18** | 6 - 28 | 15 | 3.19** | 15 - 37 |
|  | D | 5 | 95 | 4.78** | 79 -105 | 102 | 5.12** | 88 - 105 |
|  | D | 16 | 6 | 4.35** | 6 - 30 | 15 | 4.43** | 15 - 49 |
| *Climbing* |  |  |  |  |  |  |  |  |
|  | A | 1 | 99 | 4.30* | 90 - 106 | 98 | 4.17* | 91 - 103 |
|  | A | 2 | 58 | 5.68* | 54 - 64 | 78 | 5.69* | 74 - 82 |
|  | A | 3 | 34 | 2.89** | 18 - 54 | 42 | 2.97** | 16 - 60 |
|  | A | 6 | 1 | 2.72** | 1 - 23 | 7 | 2.71** | 7 - 71 |
|  | A | 9 | 76 | 2.29** | 6 - 76 | 74 | 2.29** | 7 - 74 |
|  | B | 1 | 99 | 4.86* | 88 - 106 | 95 | 4.76* | 89 - 103 |
|  | B | 2 | 58 | 6.12* | 52 - 64 | 80 | 6.23* | 74 - 84 |
|  | B | 3 | 34 | 3.37** | 18 - 50 | 42 | 3.36** | 20 - 58 |
|  | B | 6 | 1 | 3.06** | 1 - 22 | 7 | 3.02** | 7 - 69 |
|  | B | 9 | 76 | 3.08** | 34 - 76 | 74 | 3.02** | 27 - 74 |
|  | B | 13 | 10 | 3.32** | 8 - 16 | 26 | 3.17** | 26 - 32 |
|  | B | 15 | 2 | 3.90** | 2 - 8 | 6 | 3.77** | 6 - 12 |
|  | B | 16 | 24 | 5.36* | 14 - 28 | 20 | 5.14* | 19 - 43 |
|  | B | 17 | 42 | 3.95** | 35 - 42 | 58 | 3.84** | 45 - 58 |
|  | C | 1 | 99 | 5.31* | 92 - 102 | 98 | 5.21* | 93 - 101 |
|  | C | 2 | 58 | 5.80* | 52 - 64 | 78 | 5.79* | 74 - 84 |
|  | C | 3 | 34 | 4.34** | 20 - 46 | 42 | 4.44* | 26 - 50 |
|  | C | 16 | 22 | 3.17** | 12 - 26 | 21 | 3.08** | 19 - 43 |
|  | D | 1 | 99 | 5.86* | 92 - 104 | 98 | 5.75* | 93 - 103 |
|  | D | 2 | 58 | 6.23* | 52 - 64 | 80 | 6.35* | 74 - 84 |
|  | D | 3 | 34 | 4.83** | 20 - 44 | 42 | 4.87** | 28 - 52 |
|  | D | 15 | 2 | 4.02** | 2 - 10 | 6 | 3.88** | 6 - 64 |
|  | D | 16 | 22 | 6.82* | 16 - 26 | 20 | 6.72* | 19 - 35 |
|  | D | 17 | 42 | 4.20** | 35 - 42 | 58 | 4.13** | 45 - 58 |

\*Significant QTL exceeding the 0.05 genomewide adjusted threshold
\*\*Suggestive QTL exceeding the 0.63 genomewide adjusted threshold

### The impact of the revised genetic map on QTL localization

Reanalyzing QTL using the refined rat genetic map revealed striking differences to localization and significance of multiple previously mapped loci (Table 2). As we did not detect disordered markers in the revised map compared with the original dataset (SOLBERG et al. 2004), the QTL changes were likely due to the refinement of genetic distance between markers and its impact as a function of interval mapping (LANDER and BOTSTEIN 1989). Our findings recapitulate similar changes in QTL peaks that were observed following revision of the mouse genetic map (COX et al. 2009) and demonstrate the importance of a high-resolution genetic map for QTL localization.

### Conclusions

In summary, this study has provided a substantial refinement to the rat genetic map, which had not been updated since 2004 (JENSEN-SEAMAN et al. 2004). The refined rat genetic map provides sub-centimorgan resolution and sex-specific maps for the first time, which highlighted the dramatic variation in recombination rates between males and females. An added advantage of the revised rat genetic map is its derivation from the eight common inbred founder strains within the HS rat population and therefore the large number of genetic loci that were placed on the map are likely to be applicable to the majority of independent linkage analyses. Importantly, the refined marker positions in the sex-averaged map strongly impacted the localization, shape, and effect size of multiple QTL, indicating that other existing and future QTL mapping studies might be similarly affected. The refined rat genetic map will also serve as a significant resource to the growing community of geneticists utilizing outbred rat models to fine-mapping QTL, such as the HS rat (KEELE et al. 2018; RAT GENOME et al. 2013; SOLBERG WOODS 2014). As such, we have provided for the rat research community, the refined genetic distances for all physical positions in the RGSC 6.0/rn6 rat genome build (1kb bins; File S2), for all genotypes on the RATDIV genotyping array (File S3), and for all SSLP markers with physical positions in the RGSC 6.0/rn6 rat genome build (File S4). These data represent an unprecedented genetic resource for mapping candidates in the rat and are on par with the current genetic map resources that are currently available for the human and mouse.

## ACKNOWLEDGEMENTS

This work was supported by grants from the NCI (R01CA193343), the Mary Kay Foundation (Grant No. 024.16), the Wisconsin Breast Cancer Showhouse Foundation, and the Advancing a Healthier Wisconsin Endowment to MJF. Additional support was received from the Wellcome Trust (089269/Z/09/Z) to AB and the NIDDK (R01DK088975, R01DK106386) and the Individualized Medicine Institute at the Medical College of Wisconsin to LCSW.

